# Deep Learning Identifies Cardiomyocyte Nuclei in Murine Tissue with High Precision

**DOI:** 10.1101/2020.01.10.900936

**Authors:** Shah R. Ali, Dan Nguyen, Brandon Wang, Steven Jiang, Hesham A. Sadek

**Affiliations:** Department of Internal Medicine, Division of Cardiology, The University of Texas Southwestern Medical Center, Dallas, Texas 75390, USA; Department of Radiation Oncology, Medical Artificial Intelligence and Automation Laboratory, The University of Texas Southwestern Medical Center, Dallas, Texas 75390, USA; Center for Regenerative Science and Medicine, The University of Texas Southwestern Medical Center, Dallas, Texas 75390, USA

## Abstract

Proper identification and annotation of cells in mammalian tissues is of paramount importance to biological research. Various approaches are currently used to identify and label cell types of interest in complex tissues. In this report, we generated an artificial intelligence (AI) deep learning model that uses image segmentation to predict cardiomyocyte nuclei in mouse heart sections without a specific cardiomyocyte nuclear label. This tool can annotate cardiomyocytes highly sensitively and specifically (AUC 0.94) using only cardiomyocyte structural protein immunostaining and a global nuclear stain. We speculate that our method is generalizable to other tissues to annotate specific cell types and organelles in a label-free way.

---

Proper identification and annotation of cells in mammalian tissues is of paramount importance to biological research. Various approaches are currently used to identify and label cell types of interest in complex tissues. In the heart, for example, immunostaining using cellspecific antibodies can be used to identify cardiomyocytes amidst vascular cells, immune cells, and fibroblasts. However, immunostaining can have limitations due to the high autofluorescence of certain tissues, nonspecific antibody binding, and/or epitope non-specificity.^1^ Another technical issue for cardiomyocytes is their large size, complex morphology, and multinucleation, which makes identification of their nuclei a complex process that requires training and the application of rigorous techniques. Genetic fluorescent reporter models can provide the most incontrovertible labeling of a cell type, but they require specialized transgenic mice.^2^ In the field of mammalian cardiac regeneration, lack of precise cardiomyocyte nuclear markers is likely one source of controversy over the wide range of reported cardiomyocyte renewal rates: misidentification of a dividing nonmyocyte nucleus can lead to calculation of a falsely high myocyte mitotic index.^1-3^

Label-free techniques have been increasingly utilized in a wide array of microscopy and biological applications.^4^ Therefore, we sought to develop an artificial intelligence (AI) tool that can identify cardiomyocyte nuclei in mouse heart tissues. To train an AI to discern cardiomyocyte nuclei from nonmyocyte nuclei, we provided a global nuclear stain (DAPI) image, a cardiomyocyte structural protein immunostained image (troponin T), and a “ground truth” (GT) cardiomyocyte nucleus-labeled image by which the AI could iteratively learn (Fig. 1A) – all from 8 μm-thick, frozen sections. GT cardiomyocyte nuclei labeling was achieved by immunostaining Myh6-Cre transgenic mice cardaic tissue for Cre (red) (Fig. 1E); the labeled cardiomyocyte nuclei are readily distinguished from nonmyocyte nuclei. However, we also noted that there is extranuclear Cre tissue staining, so we subtracted the tissue image (Troponin stain) from the Cre-stained image (Fig. 1F,G): the resulting processed image was treated as the input GT label for training the neural network.

**Fig 1:**
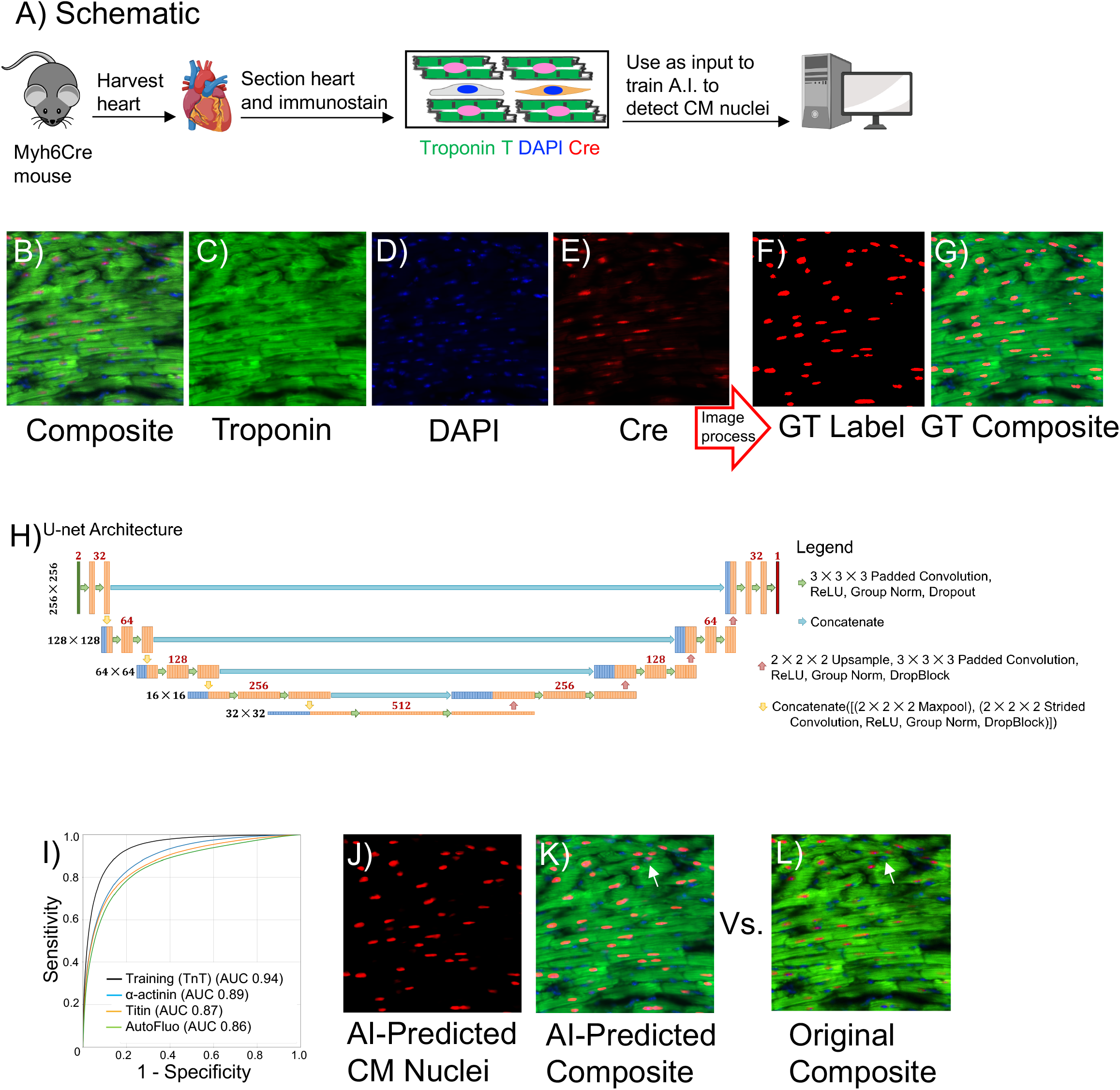
A) Schematic of overall approach. B-E) Immunostained images of Myh6Cre frozen heart sections (8 μm-thick) using Troponin (Millipore 06-570, 1:100) and Cre (Millipore 69050, 1:10000) antibodies. F-G) Processing of the Cre-immunostained image renders a Ground Truth (GT) label that is used to train the AI model H) Schematic of the U-net architecture used in this study. The black numbers at the left represent the image resolution at each level of the U-net. The red numbers represent the number of feature maps that are calculated when the convolution is applied. I) Receiver-operating curve (ROC) of the test model (troponin T, black) and alternate cardiomyocyte structural immunostained-images. Standard deviations: Troponin T 0.032, α-actinin 0.004, titin 0.027, and autofluorescence 0.019 (n=3 mice per stain). J-L) Image of AI-predicted cardiomyocyte (CM) nuclei and original GT CM labels; arrowhead in (K, L) points to nonmyocyte nucleus mislabeled by the AI as a myocyte nucleus).

Since all of the data inputs and labels are on the pixel level we used an image-to-image deep learning model, a U-net style architecture^5^, which is one of the most widely used styles of architectures for imaging tasks that can calculate and maintain both global and local information throughout the network (Fig. 1H).

To evaluate the model’s stability and robustness we trained 5 different models, but withheld a portion of the training data to use it as validation data. We divided each complete heart section into 256 x 256 patches. For the 25 tissue sections used as training and validation data (n=3 mice), this resulted in 33,769 total image patches for model development. For each model, 27,016 image patches were used for training and 6,753 were left out and used for validation.

Each instance of the model was trained for a total of 200,000 iterations with a batch size of 4, which took approximately 20 hours 36 minutes (standard deviation 47 minutes), on an NVIDIA 1080 Ti GPU. For each training instance, the best performing weights with the lowest validation score was used. All 5 model instances took 4.29 days to train for 200,000 iterations each. All 5 models were evaluated on the testing set, to obtain the model’s overall performance and variation.

All models performed well: the models’ prediction closely matched the ground truth label and had a bright signal where they matched (Fig. 1J-L); when the model mislabeled a nonmyocyte as a myocyte, its assigned signal is dimmer (arrowhead, Fig. 1K). The average area under the curve (AUC) across all 5 models was 0.940 ± 0.032 (Fig. 1L, black curve), which demonstrates that the models have a high probability of accurately distinguishing a nonmyocyte from a myocyte nucleus (AUC 1 would indicate a perfect model). The model’s accuracy decreased only slightly when tested upon images with alternative structural protein stains (α- actinin, titin) or no stain (autofluorescence) (Fig. 1L), validating the model’s reliance on the cardiomyocyte sarcomeric protein pattern for accuracy.

Misidentification of cells in biological research can confound interpretation of underlying physiological processes. Herein, we demonstrate that machine learning-based methods may automate identification of an important cardiac cell type. We generated an AI model that uses image segmentation to predict cardiomyocyte nuclei in a mouse heart section without a specific cardiomyocyte nuclear label. This tool, which can annotate cardiomyocytes highly sensitively and specifically (AUC 0.94) using only pan-nuclear and cardiomyocyte structural protein immunostains, enables calculations that were previously inaccessible to a manual approach (e.g., counting all the cardiomyocyte nuclei in a tissue section). More experiments, however, are needed to determine the applicability of this model to tissue sections of different thickness and orientation and to pathological tissues. We speculate that our approach is generalizable to other organs to annotate specific cell types and organelles in a label-free way.

## APPENDIX

### Tissue Processing and Staining

Tissue from 8-week-old Myh6-Cre mice were fixed in 4% paraformaldehyde overnight at 4°C, cryopreserved in 30% sucrose at 4°C, and embedded in Tissue Freezing Medium (TFM). The cryomolds were sectioned at 8 μm thickness (Leica cryostat), followed by antigen retrieval (by boiling the slides in 1 mM EDTA/0.05% tween for 40 minutes), and immunostained for Cre protein (rabbit 1:10000, Millipore Sigma #69050), troponin T (mouse 1:100, Millipore 06-570), and counter-stained with DAPI (Thermo-Fisher). Alternate structural protein stains were performed in the same way but using α-actinin (mouse 1:100, Abcam ab18061) or titin (mouse 1:100) or none (autofluorescence). Images were taken from these immunostained sections at 20x. There are 3 channels for each image: the tissue stain (troponin T, green), the nuclei stain (DAPI, blue), and the cardiomyocyte nuclear stain (Cre, red) (Fig. 1B-G).

### Training Data Preparations

We used the tissue and nuclei stain as the input to teach the deep learning model to predict the cardiomyocyte nuclei. To achieve this, it is required to convert the cardiomyocyte nuclei stain into labels to teach the model. However, since the cardiomyocyte stain still contains a lot of tissue information, we executed multiple steps in order to create the proper image of cardiomyocyte labels, which is outlined below.

The first step is to remove as much of the tissue information as possible, to achieve this we subtracted the tissue image from the cardiomyocyte image,

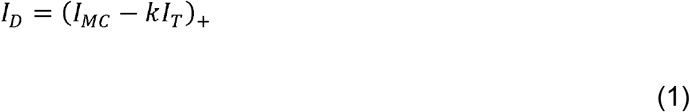

where *I_MC_* is the cardiomyocyte image, *I_T_* is the tissue image, *I_D_* is the resulting difference image, and k is a scalar constant. The operator, (·)_+_, projects its argument onto the nonnegative orthant. With the goal of removing as much tissue information, we optimized k for each slide image, based on the minimization problem:

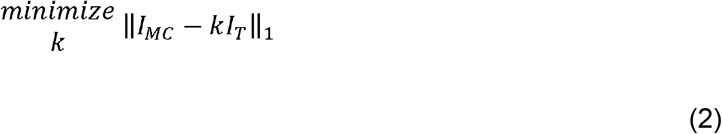

where ‖·‖_1_ is the *l*_1_-norm. In mathematics, the *l*_p_-norm of an argument, 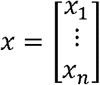, is defined as 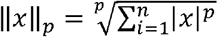. We chose the *l*_1_-norm as our fidelity term, since the *l*_1_-norm promotes its argument to be sparse in a minimization problem. Since the /_r_norm is non-differentiable at 0, we utilized a first-order primal-dual proximal-class algorithm known as Chambolle-Pock^6,7^, which can handle non-differentiable functions through the use of the prox operator, which is defined, for some function, *h*(·), and a step size, *t*, as 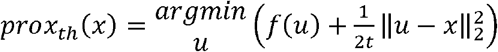. After solving Equation 2 to find k, we can apply Equation 1 to obtain the difference image, *I_D_*. The reason why we project the difference onto the non-negative orthant is due to the fact that the cardiomyocytes nuclei have higher fluorescence intensity than the surrounding tissue in the Cre immunostain. Therefore, we expect that all cardiomyocyte nuclei will be positive after the operation *I_MC_ − kI_T_*. Finally, to create the labels, we must first separate the cardiomyocytes from any residual background in *I_D_*. To do this, we used a method known as Otsu thresholding^8^, which is an advanced thresholding technique to separate an image into a foreground and background. It automatically selects the threshold based off an intensity histogram, minimizing the variance within a class and maximizing it between the two classes. We applied this method onto *I_D_* to arrive at our final cardiomyocyte label image, *I_MCL_*, shown in Equation 3:

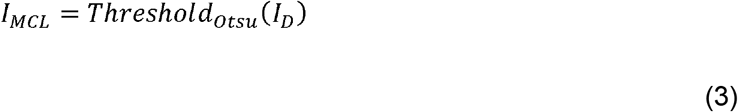

This *I_MCL_* is then treated as the ground truth label for training the neural network to learn cardiomyocyte nuclei identification.

### Deep learning architecture

Since all of the data inputs and labels are on the on the pixel-level, using an image-to-image deep learning model is the natural choice for this study. We utilized a U-net style architecture^5^. The U-net architecture was proposed in 2015 and was originally applied for biomedical image segmentation tasks. The strength of the U-net comes from its capability to calculate and maintain both global and local information throughout the network, and has become one of the most widely used styles of architectures for imaging tasks. Our exact variant of the U-net is shown in Figure 1H. The basic block of operations used in this study involves a convolutional layer, a rectified linear unit (ReLU) as the non-linear activation, group normalization^9^ (GN), and DropBlock^10^ (DB). Group normalization was developed in 2018 to overcome the limitations of batch normalization^11^, especially with small batch sizes. DropBlock is a special variant of dropout^12^, and was shown to work better with convolutional neural networks than the standard dropout. We will refer to this block of operations as Conv-ReLU-GN-DB.

To be specific, our network was developed to take in 2 channels of 256 x 256 image patches and output 1 channel of the same resolution. A total of 5 resolution levels were designed, with downsampling and upsampling operations in between to translate the data between different resolutions. The downsampling consists of 2 parallel operations, whose outputs are then concatenated together. The first operation is a 2 x 2 max pooling^13,14^, and second set of operations is a 2 x 2 strided Conv-ReLU-GN-DB with a 2 x 2 kernel. For the upsampling, we performed a nearest neighbor 2 x 2 upsample, followed by a Conv-ReLU-GN-DB operation with a 3 x 3 kernel. Between these downsampling and upsampling operations, 2 sets of Conv-ReLU-GN-DB with a 3 x 3 kernel are performed. Regarding the number of feature maps calculated at each convolution, the first level starts at calculating 32 feature maps at each layer, the lower levels of decreasing resolution double the number of feature maps to 64, 128, and 256, respectively. The model development and training framework was developed using TensorFlow^15^. The Adam optimizer^16^ was used during training, with a learning rate of 1 × 10^−3^.

### Training and Evaluation

To evaluate the model’s stability and robustness, we trained 5 different models, omitting a different portion of the training data from training, which was then used as validation data. We divided each slide image into 256 x 256 patches. For the 25 slides used as training and validation data, this resulted in 33,769 total image patches for model development. For each model, 27016 image patches were used for training and 6753 were left out and used for validation. For each training instance, the weights where the model had performed best on the validation set was then used to evaluate on the testing data, in order to assess the average prediction performance and variation.

In total, on the 2 slides used as hold-out testing data, there are 81,330 nuclei in the testing data that was evaluated. To determine whether the model’s prediction should be labeled as cardiomyocyte or not, we define the fraction of voxels labeled as positive in the *i^th^* nuclei, *f_i_* as

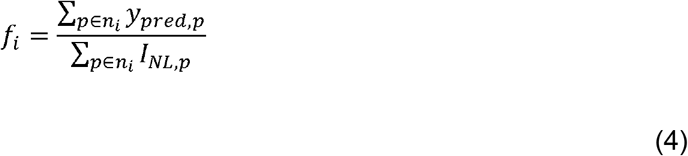

where *p* is the pixel index, *n_i_* is the *i^th^* nuclei, *y_pred_* is the model’s raw prediction of cardiomyocytes, and *I_NL_* is the image of nuclei labels, obtained by performing Otsu thresholding on the nuclei stain image. The summation over *p* ∈ *n_i_* means that we are only summing over the voxels defined in the *i^th^* nuclei. We can then define the nuclei as cardiomyocyte if *v_fi_* exceeds a defined threshold value

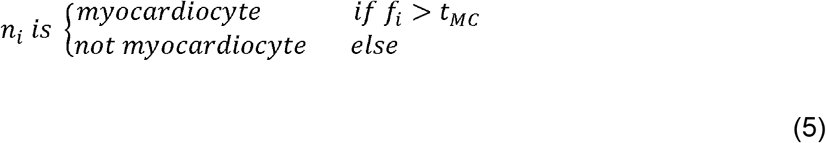

where *t_MC_* ∈ [0,1] is a user defined threshold, and controls the tradeoff between the sensitivity and specificity. We calculated the true/false positives/negatives for each *t_MC_* between 0 and 1, with a step size of 0.001. In total, we evaluated the prediction using 1000 *t_MC_* values.

